# An AI/ML-Powered Workflow for End-to-End Cell Line Development

**DOI:** 10.64898/2026.02.04.703387

**Authors:** Shalini Raj Unnikandam Veettil, Joanna Donatelli, Geetansh Kalra, Constanza Verónica Ljubetic San Martín, Shankar Ramakrishnan, Collin McGregor, Michelle Wallace, Ramya Ankala, Leonardo Rodrigues de Souza Pinto, Arun Dhama, Chris Regens, Yi Li, Derek K. Smith

**Affiliations:** Gilead Sciences, Foster City CA; E Tech Group, West Chester OH; Thoughtworks Inc., Chicago IL

## Abstract

The generation of clonal CHO cell lines is foundational to biologics manufacturing; however, labor-intensive cell culture workflows predominate in the field. We created the CLAIRE (Cell Line AI Recognition and Evaluation) tool to streamline end-to-end cell line development by integrating deep-learning image analysis with automated liquid handling. We benchmarked three object detection models for monoclonality verification and found DETR provides superior accuracy (>0.90 F1-score) in identifying single cells. To quantify the outgrowth of cell lines, we evaluated multiple zero-shot SAM2 segmentation models against a feature-based estimation method. Feature-based detection successfully identified diverse cell colony types while less robust performance was observed for SAM2 models, particularly for sparse density colonies. The pre-trained DETR and feature-based detection models were wrapped in a task-focused user interface that outputs cell line hitpick lists compatible with a Lynx LM1800 liquid handler in addition to custom scripts automating cell passaging and sampling. This approach yielded an end-to-end 36 day CLD workflow capable of generating high-titer cell lines for multiple complex antibody structures. Here, we open-access our trained models, user interface, and Lynx automation scripts to provide a modular toolkit useful for clonal cell line engineering projects.

**Graphical Abstract:** 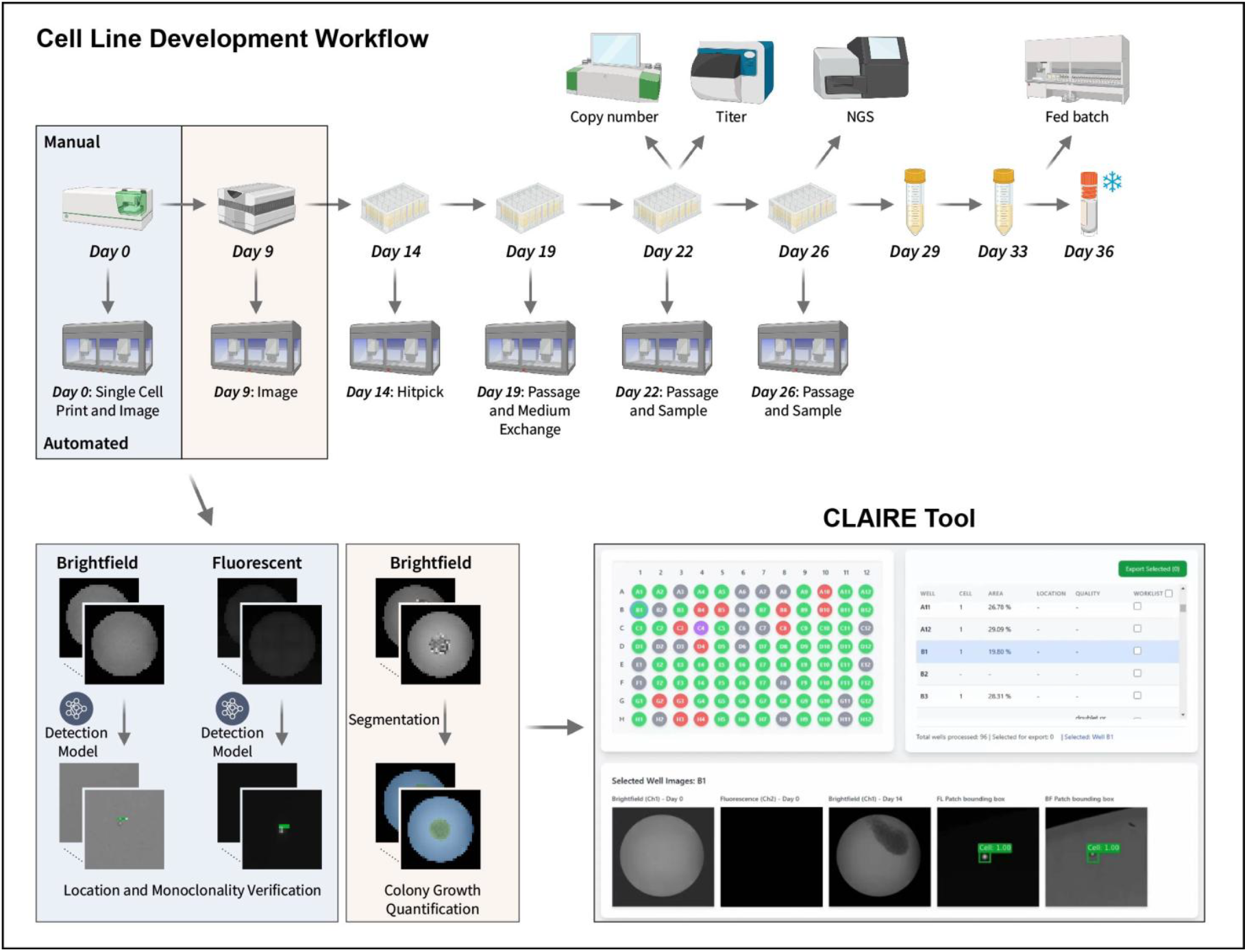

## Introduction

Industry-leading cell line development (CLD) platforms engineer Chinese Hamster Ovary (CHO) cells to generate stable, high-yield clonal cell lines for recombinant protein expression. While effective for large-scale manufacturing, these cell lines can exhibit heterogeneity in growth, protein yield, and product quality. For this reason, regulatory bodies require that biologics production utilize monoclonal cell lines with proof of single cell origin from image-based documentation or confirmatory statistical methods (1-3).

The identification of high-producing clonal cell lines is a time- and labor-intensive process. While state-of-the-art cell sorters enable high-throughput single cell deposition coupled with nozzle imaging (4), independent image-based confirmation of monoclonality in the destination well is beneficial to regulatory filings. Even accurate sorters such as UP.SIGHT (CYTENA) are prone to infrequent deposition error with 0.29% of wells containing cell doublets or multiples (5). In cases where post-deposition imaging is performed, accurate assessment of well images can be complicated by atypical cell morphology or image artifacts such as shadows, debris, or well-edge curvature effects. These variables make manual image review prone to human error and inter-operator variability.

To overcome these challenges and accelerate development timelines, biopharmaceutical frontrunners have integrated automated liquid handling and digitization into CLD workflows (6). This shift increasingly leverages computer vision to apply deep learning and statistical methods to image classification, object detection, spatial localization, and tracking. Convolutional neural networks (CNNs) automatically detect patterns using layered filters and eliminate the need for manual feature extraction (7). General-purpose tools in the NIH ImageJ ecosystem (8-12) such as DeepImageJ (13) and CellProfiler (14) enable the use of deep-learning models and automate the identification of pre-trained cell phenotypes, but neither tool is optimized to output large-scale structured data for automated CLD workflows. Specialized tools facilitate high-precision cell segmentation (15-18) and enumeration (19, 20) such as StarDist (21) and AICellCounter (22), a light-weight AI-based solution for cell counting. Despite these advancements in general and specialized image processing, none sufficiently address our need for monoclonality verification and confluency measurements coupled to automation-friendly hitpick list generation in a user-focused graphical interface.

In this article, we open-access an end-to-end CLD process including Lynx LM1800 scripts to automate cell culture and the image analysis deep learning framework with user interface CLAIRE (Cell Line AI Recognition and Evaluation). We compared three state-of-the-art object detection algorithms (DETR, Faster R-CNN ResNet50 FPN-2 (FRRF-v2), and YOLO-NAS) and benchmarked performance for monoclonality, doublet, and multiple cell detection. Further, we evaluated two methods (SAM2 and image feature-based estimation) for estimating colony confluency.

We developed a single-click user interface that utilizes the outputs of these analyses to generate formatted hitpick lists compatible with a Lynx LM1800 liquid handler (Supplemental Data). Lynx LM1800 liquid handlers enable cell count-mediated passaging and normalization through volume-verified pipetting and simultaneous transfer of 96 unique volumes from source to destination wells. We created time-optimized scripts to enable dynamic clone hitpick, cell passaging, scale-up, and sample collection for ddPCR, next-generation sequencing, and titer assays. We present data from cell lines generated using this workflow to demonstrate process robustness and cell line performance across a variety of antibody formats.

To encourage adoption, we open-access the trained models, analysis scripts, installable CLAIRE interface, cell culture workflow, and Lynx LM1800 method scripts. This comprehensive, open-access toolset is designed to simplify scientist operations by automating image ingestion, image analysis for monoclonality and confluence, hitpick list generation, and cell culture workflows for CLD.

## Results

### Dataset Generation for Model Training

Training datasets were generated by printing individual CHOZN GS^-/-^ CHO cells into the wells of 40 Greiner 96-well plates (655090) and 40 Corning 96-well plates (3596) followed by same-day brightfield and fluorescent imaging of each well. Cells were incubated for 14 days and brightfield imaging of each well performed to capture cell colony outgrowth (**Figure 1a**). Training on full 7530 × 7530 pixel images was inefficient because each single cell occupied less than 0.0004% of the area (**Figure 1b**). To address this, smaller regions of interest of 256 × 256 pixels were extracted from the focus control well of each plate which contained thousands of cells that were added to enable focal plane detection prior to imaging. Utilizing focus control wells enabled training not only on single cells, but more complex doublet and cluster structures. This resulted in 1,500 brightfield and fluorescence images annotated as single cells, doublets, and clusters. (**Figure 1c-f**).

**Figure 1.**
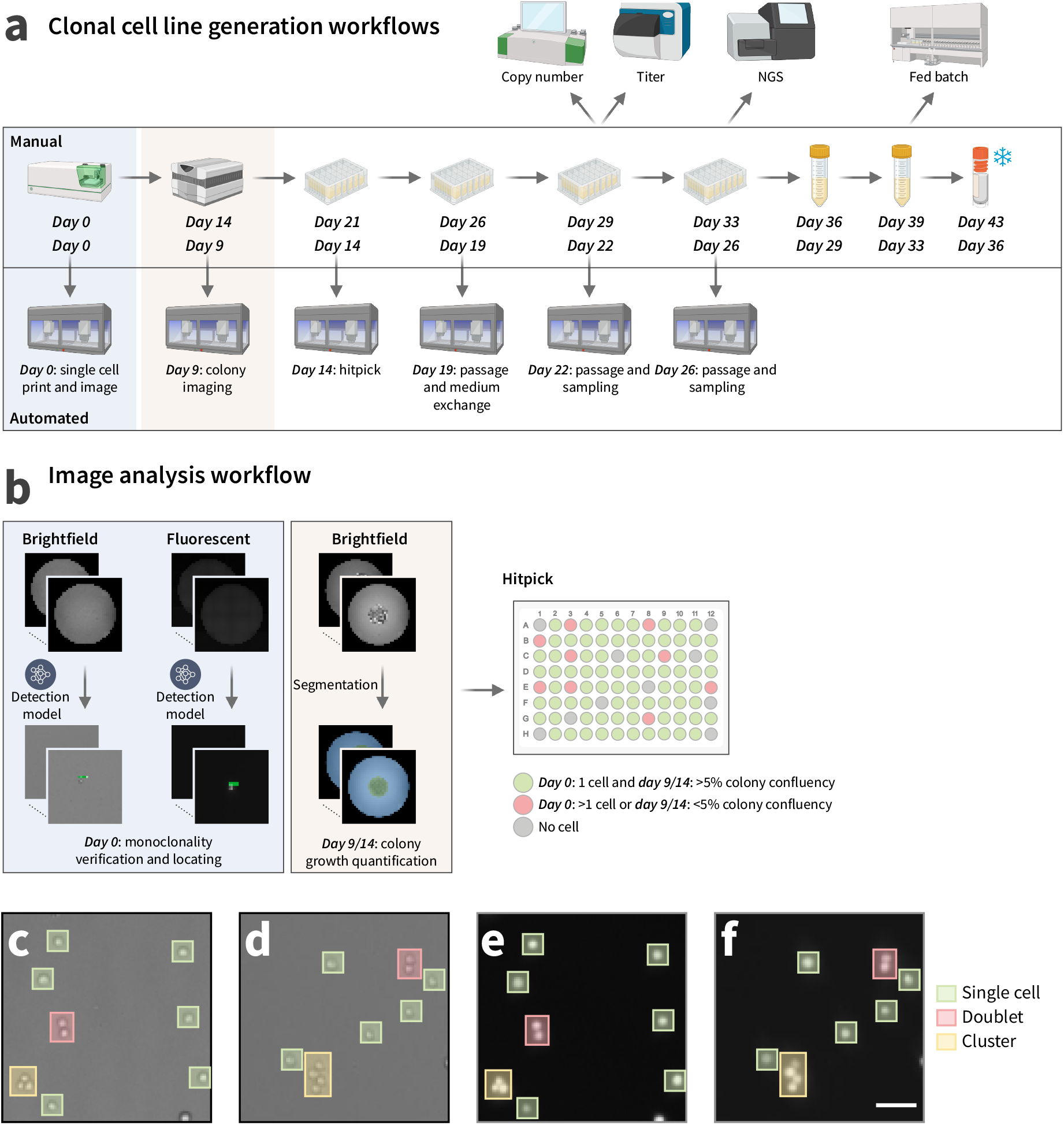
(a) Clonal cell line generation workflow. (b) Image analysis workflow for brightfield and fluorescent imaging. Detection models verify day 0 monoclonality and segmentation quantifies day 9/14 colony growth. Hitpick is determined from monoclonality and colony confluency thresholds. (c-f) Representative well images with bounding boxes for annotation classes: single cells, doublets, and cell clusters. Brightfield images for (c) Greiner and (d) Corning plates. Fluorescent images for (e) Greiner and (f) Corning plates. Scale bar: 50 μm.

Model performance was evaluated using two test sets. The first consisted of image patches from focus control wells seeded at high density, which include cell doublets and clusters, to test the accuracy of each model’s monoclonality prediction. The second consisted of image patches from wells where only a single cell was deposited to represent the standard workflow. Predictions were evaluated using an Intersection over Union (IoU) threshold of 0.5 for bounding box matching and a confidence threshold of 0.7 for accepting detections.

### Performance Evaluation: High Cell Density Images Similar to Training Datasets

DETR delivered the strongest overall performance across both datasets among the evaluated models. DETR achieved high overall F1-scores (0.95 brightfield, 0.79 fluorescent), precision scores (0.95 brightfield, 0.78 fluorescent), and recall scores (0.95 brightfield, 0.87 fluorescent) (**Figure 2a, b**). While FRRF-v2 and YOLO-NAS performed comparably for brightfield images with overall F1-scores (0.89 FRRF-v2, 0.85 YOLO-NAS), YOLO-NAS recorded the lowest overall F1-score (0.75) for fluorescent images. Overall model performance was higher for brightfield images relative to fluorescent images (**Table 1**). This performance is particularly relevant to the single cell deposition workflow since false doublet classification directly impacts regulatory confidence.

**Table 1.** Precision, recall, and F1-score for FRRF-v2, YOLO-NAS, and DETR models evaluated on brightfield and fluorescent test images. Metrics are reported for overall performance across all classes and for individual object classes: single cell, doublet, and cluster.

|  |  | Brightfield |  |  |  | Fluorescent |  |  |  |
| --- | --- | --- | --- | --- | --- | --- | --- | --- | --- |
| Model | Metric | Overall | Single cell | Doublet | Cluster | Overall | Single cell | Doublet | Cluster |
| FRRF-v2 | Precision | 0.94 | 0.98 | 0.93 | 0.95 | 0.78 | 0.92 | 0.92 | 0.49 |
|  | Recall | 0.84 | 0.86 | 0.82 | 0.85 | 0.83 | 0.92 | 0.69 | 0.89 |
|  | F1-score | 0.89 | 0.90 | 0.87 | 0.89 | 0.78 | 0.92 | 0.79 | 0.63 |
| YOLO-NAS | Precision | 0.95 | 0.96 | 0.94 | 0.94 | 0.80 | 0.95 | 0.95 | 0.52 |
|  | Recall | 0.77 | 0.76 | 0.69 | 0.85 | 0.76 | 0.76 | 0.62 | 0.90 |
|  | F1-score | 0.85 | 0.85 | 0.80 | 0.90 | 0.75 | 0.85 | 0.75 | 0.66 |
| DETR | Precision | 0.95 | 0.94 | 0.96 | 0.96 | 0.78 | 0.93 | 0.92 | 0.49 |
|  | Recall | 0.95 | 0.97 | 0.93 | 0.95 | 0.87 | 0.96 | 0.69 | 0.95 |
|  | F1-score | 0.95 | 0.96 | 0.94 | 0.96 | 0.79 | 0.94 | 0.79 | 0.65 |

**Figure 2.**
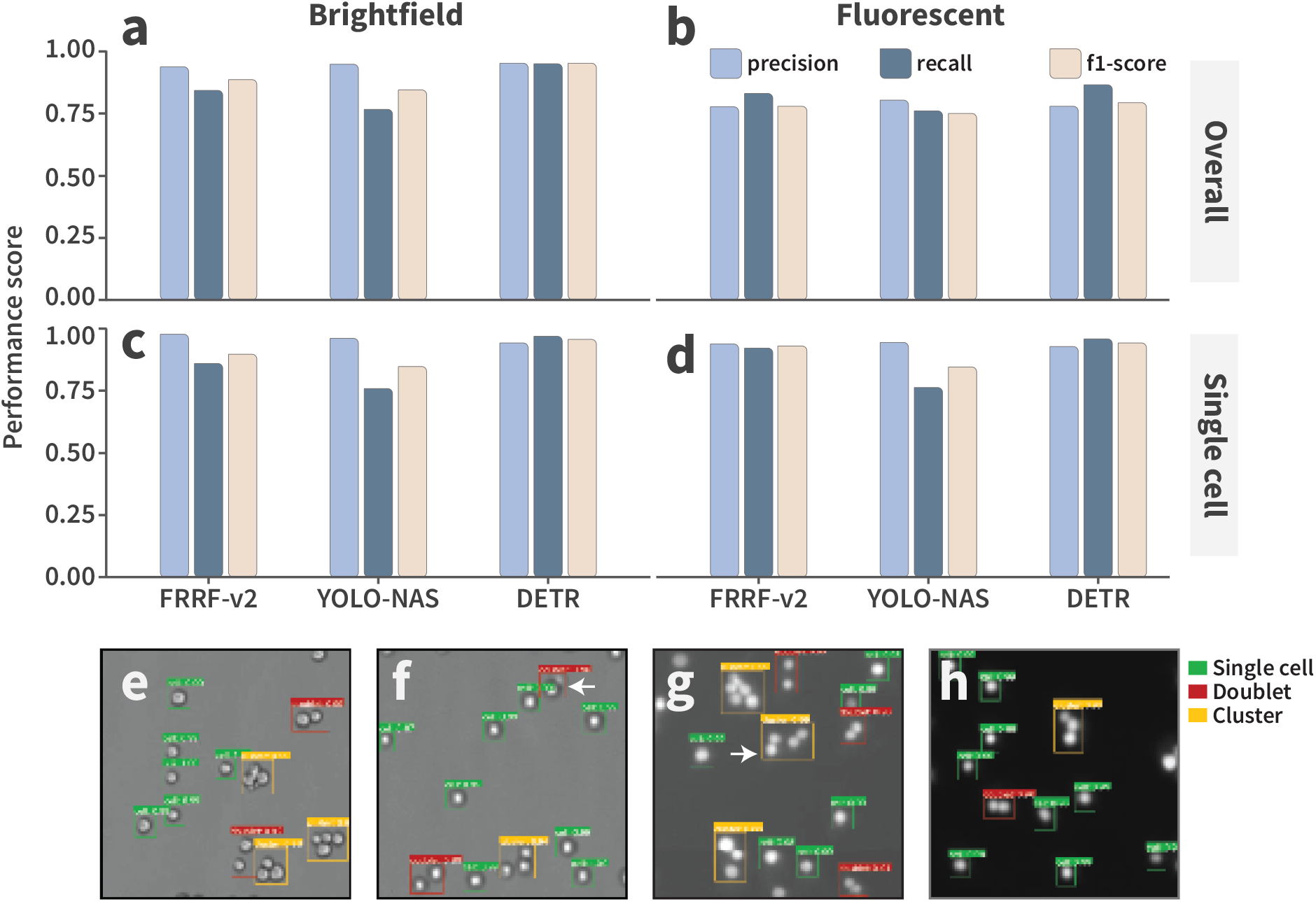
Model performance scores on test datasets. (a) Overall performance across all classes on brightfield images. (b) Performance for single-cell classification on brightfield images. (c) Overall performance across all classes on fluorescent images. (d) Performance for single-cell classification on fluorescent images. Example detections from the DETR model showing predicted bounding boxes for brightfield images from (e) Greiner plate and (f) Corning plate, as well as, fluorescent images from (g) Greiner plate and (h) Corning plate.

Although performance varied across individual classes, the *single cell* classification consistently showed highest precision, recall, and F1-scores across all models (**Figure 2c, d**). DETR had the highest F1-scores for this classification (0.95 brightfield, 0.94 fluorescent) with balanced precision and recall (0.95, 0.95 brightfield and 0.93, 0.96 fluorescent). FRRF-v2 and YOLO-NAS similarly demonstrated high precision for single cell detection in brightfield images (0.98 and 0.96, respectively); however, both yielded lower recall values (0.86 and 0.84, respectively). Precision and recall were consistent for FRRF-v2 (0.92, 0.92), while YOLO-NAS recall dropped significantly (0.76) relative to the other models.

These results show similar model performance across classifications in brightfield images and highest scores for single cell detection. In contrast, fluorescent images exhibited greater variability between classifications. Misclassification primarily occurred when doublets were grouped with adjacent cells or other doublets and predicted as a single large cluster, which led to higher false negatives for doublets and higher false positives for clusters. These issues contributed to a decline in doublet recall, which dropped for DETR (0.69), FRRF-v2 (0.69), and YOLO-NAS (0.62). For the cluster classification, precision decreased substantially for DETR (0.49), FRRF-v2 (0.49), and YOLO-NAS (0.52). In several cases, two closely adjacent doublets were not detected individually and, instead, were predicted as a single cluster (**Figure 2g**).

### Performance Evaluation: Single Cell Images from Cell Line Development Workflow

The ability of each model to distinguish cell classifications successfully, particularly single cells, prompted a second evaluation using more challenging CLD workflow images that require detection of a single cell against the sparse background of a 96-well plate. Paired brightfield and fluorescent images of individual wells were processed using a two-stage workflow. Fluorescent images were converted to grayscale, enhanced using CLAHE (clipLimit = 2.0, tileGridSize = 8,8), and binarized using adaptive Gaussian thresholding (block size: 5, constant: 2) to generate a binary mask for contour detection. Bounding boxes generated from cell contours were used to define regions of interest then corresponding brightfield and fluorescent patches of 256 × 256 pixels extracted. Images were cropped from both the center of the bounding box and randomized offsets from center. These patches were analyzed using object detection models trained specifically for brightfield and fluorescent images. Predicted bounding boxes, classification labels, and confidence scores were evaluated to determine accuracy and robustness. Patches extracted without a detectable cell, doublet, or cluster classifier were automatically flagged as unknown for manual scientist review. This capability demonstrates embedded quality control during high-throughput image processing.

Relative to training datasets, all models exhibited a decline in precision, recall, and F1-score for single cell brightfield images (**Figure 3a**). DETR maintained strong performance for fluorescent images, while YOLO-NAS and FRRF-v2 performance declined (**Figure 3b**). Of specific note, YOLO-NAS recall (0.27) declined significantly for fluorescent signal indicating poor detection sensitivity. An IoU sensitivity analysis confirmed the superior performance of DETR with >0.90 precision, recall, and F1-score for brightfield at 0.25 IoU (**Figure 3c-e**) and similar trends for fluorescent images at 0.5 IoU (**Figure 3f-h**). These results indicate that stricter IoU thresholds penalize even minor localization errors of 1 or 2 pixels due to the relatively small area of the bounding box and indicate that most detections are spatially accurate even when exact alignment is not.

**Figure 3.**
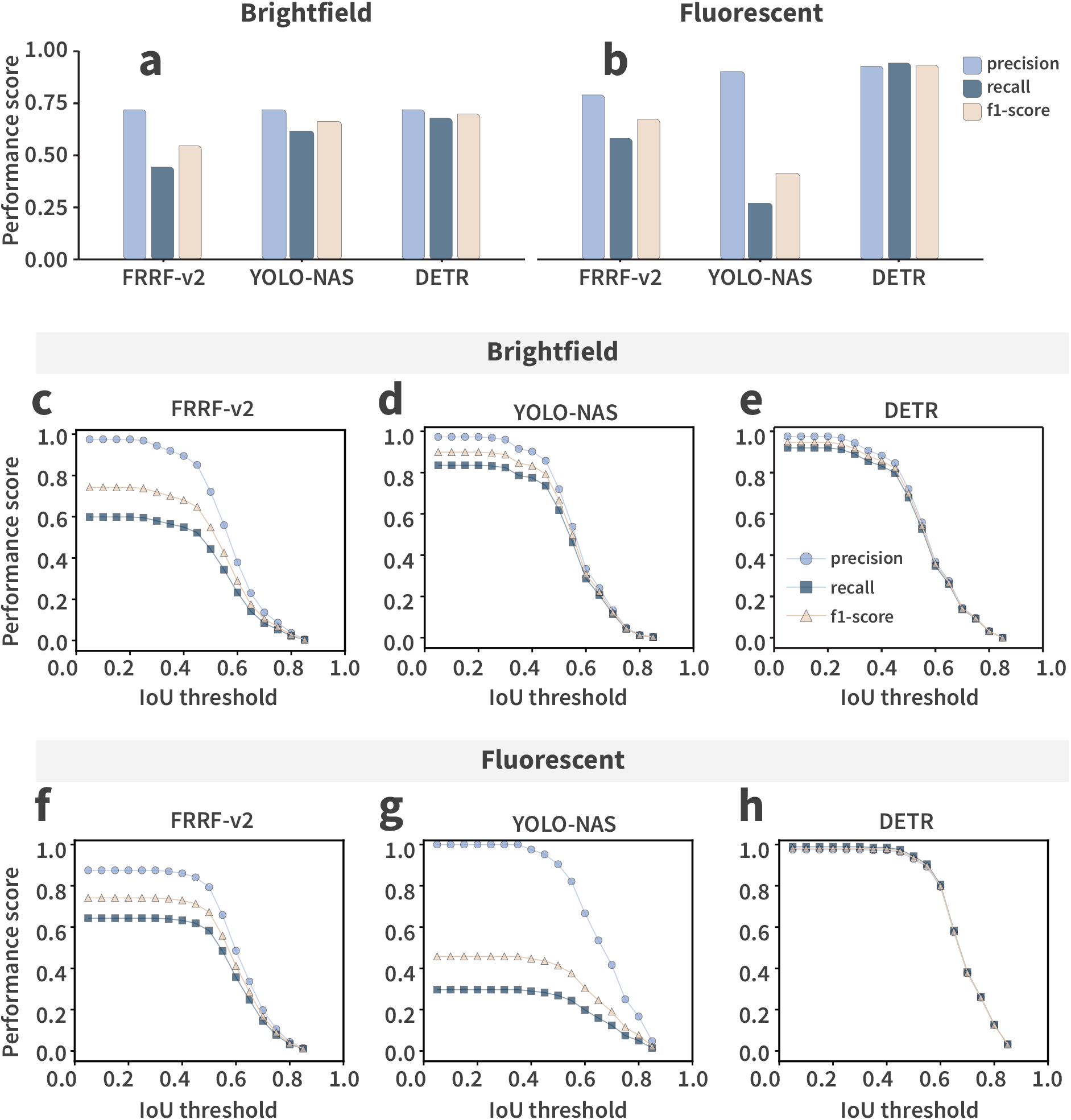
Model performance scores on test images from CLD workflow. (a) Performance for single-cell classification on brightfield images. (b) Performance for single-cell classification on fluorescent images calculated at IoU threshold of 0.5 and confidence score threshold of 0.7. (c-e) Brightfield image precision, recall, and F1-score across IoU thresholds. (f-h) Fluorescent image precision, recall, and F1-score across IoU thresholds. DETR maintains high performance at lower thresholds (<0.5) but declines sharply beyond 0.5, while FRRF-v2 and YOLO-NAS exhibit more gradual decreases.

### Performance Evaluation: Clonal Cell Line Outgrowth Quantification

Accurate segmentation of colonies with variable confluence is vital to a robust CLD workflow where novel antibody sequences could affect colony size and selection. The segmentation performance of four zero-shot SAM2 architectures with post-processing and a feature-based segmentation algorithm were evaluated for low-confluence sparse colonies, low-confluence dense colonies, high-confluence sparse colonies, and high-confluence dense colonies (**Figure 4a-f**). Brightfield images (**Figure 4a**), feature-based method segmentation outputs (**Figure 4b**), and the SAM2 hierarchical variants tiny, small, base-plus, and large (**Figure 4c-f**) are shown.

**Figure 4.**
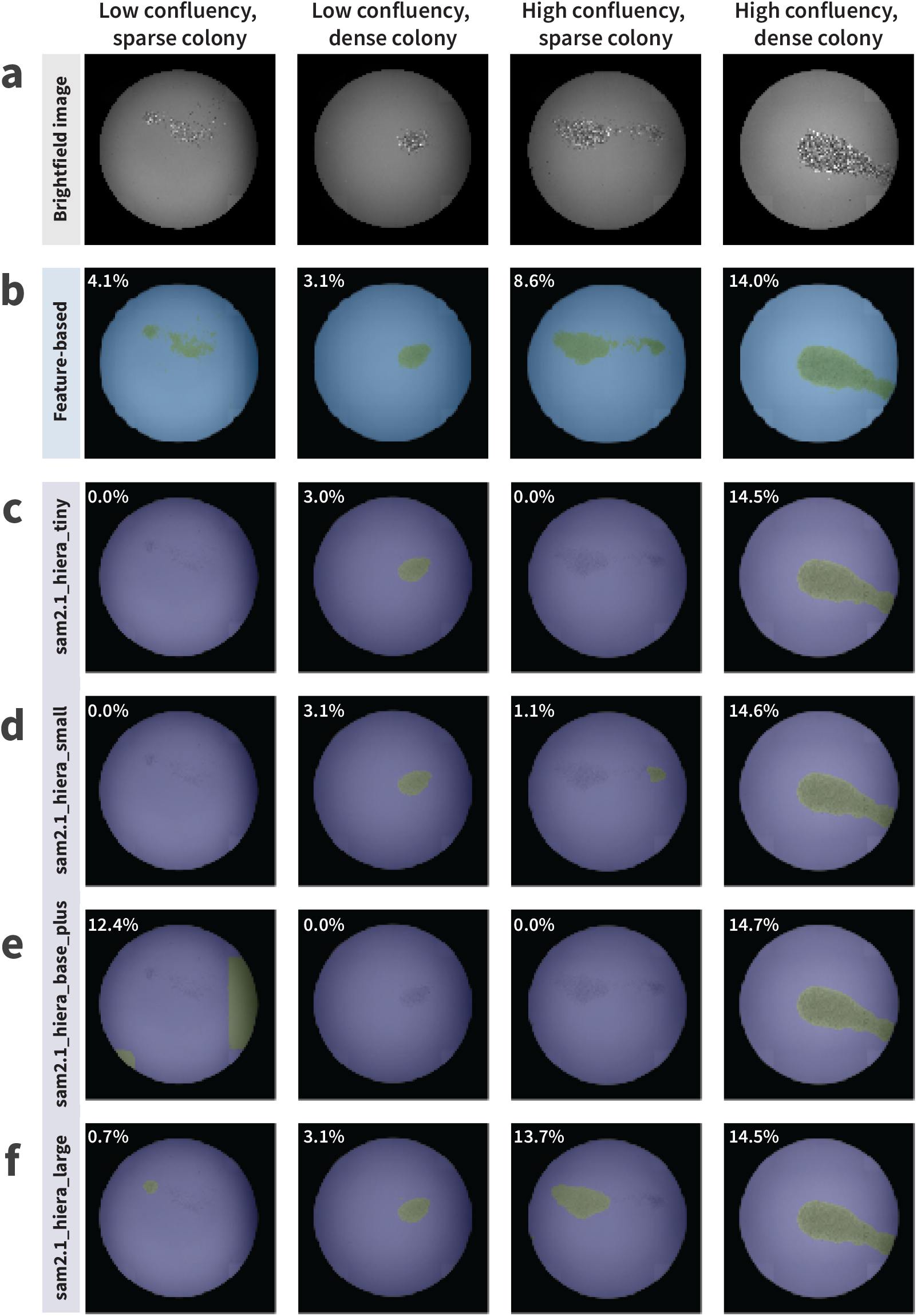
(a) Day 14 brightfield images from four representative conditions. (b) Feature-based segmentation outputs with corresponding confluency estimates (4.1%-14.0%). (c-f) Segmentation outputs from four zero-shot SAM2 architectures with post-processing. Confluency values are shown for each condition and method.

The feature-based segmentation method showed consistent performance across all cell colony classification and produced confluence estimates of 4-14% irrespective of colony morphology. In contrast, SAM2 models demonstrated strong segmentation capability for dense colonies but failed to generalize to sparse configurations often yielding null or near-zero predictions. Further, substantial variability was observed among SAM2 variants. The base-plus model occasionally over-segmented sparse colonies by incorporating imaging artifacts (**Figure 4e**). In contrast, the tiny and small models systematically under-segmented or entirely missed colonies. These results indicate that, although SAM2 architectures are effective for dense colony segmentation in zero-shot mode, their applicability to sparse conditions remains limited for this task.

### Automation of Cell Culture Workflows for Clonal Cell Line Generation

The CHOZN GS^-/-^ CHO cell line has industry-wide adoption for production of commercial antibody products. We initially subcloned this cell line using a 43-day manual workflow to determine whether genomic divergence within the host cell line could generate subpopulations with higher productivity (**Figure 5a**). Cells were stained using a fluorescent dye then 7,680 single cells were printed into 96-well plates using an F.SIGHT 2.0 single cell dispenser. Whole-well images were collected for both brightfield and fluorescent channels using a Celigo plate imager immediately following single cell deposition. Brightfield images of each well were collected after 14 days to assess colony outgrowth. Clonal cell lines were expanded and 298 cell lines were cryopreserved based on growth characteristics.

**Figure 5.**
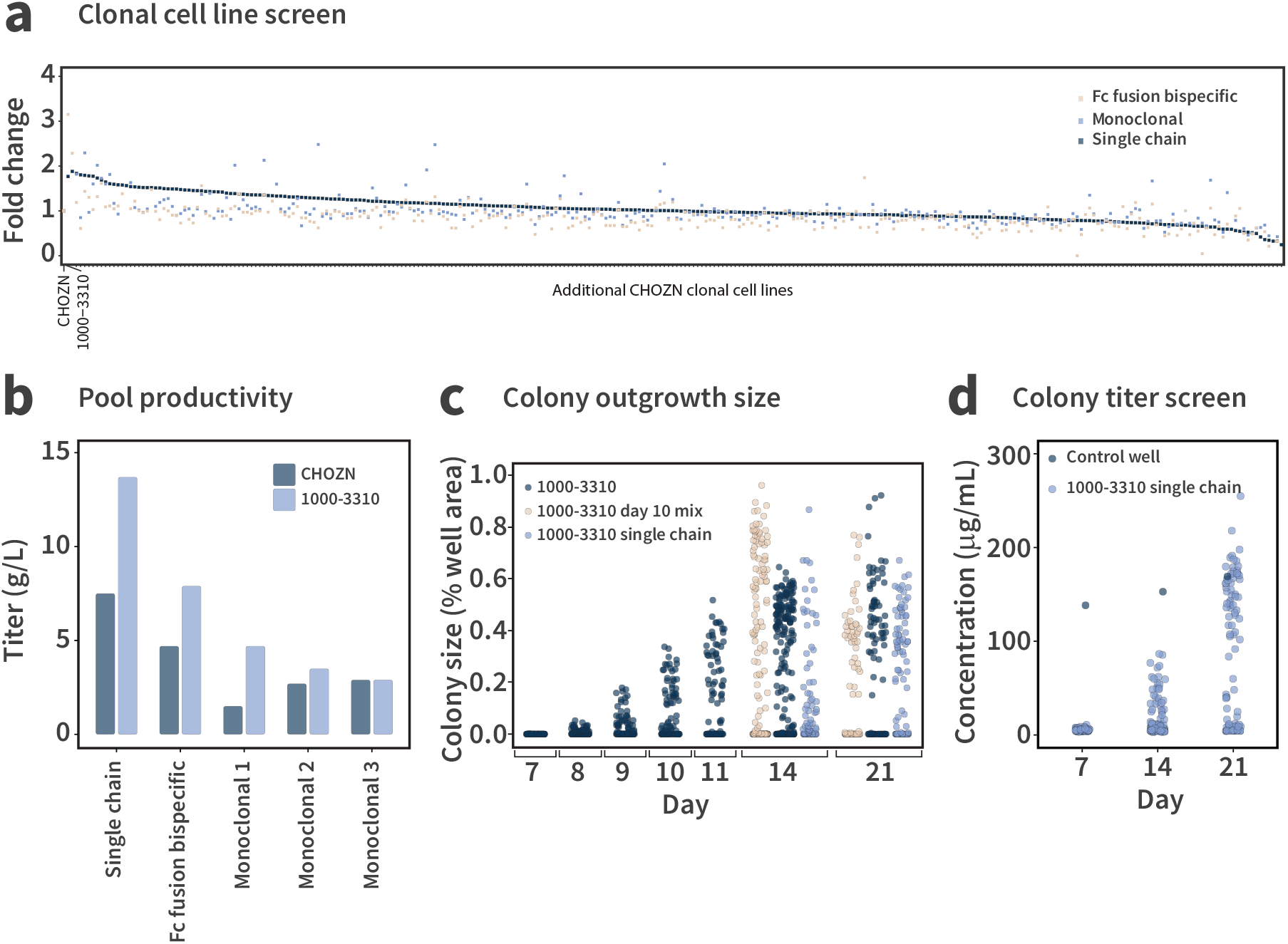
(a) The fold change (clonal cell line pool / CHOZN pool) in fed-batch titer of 298 antibody-producing clonal cell lines. (b) Fed-batch titer of CHOZN and 1000-3310 cell line pools producing five diverse antibody formats. (c) Time course of cell colony outgrowth for 1000-3310 and 1000-3310 stably expressing a single chain antibody. (d) Batch titer of clonal 1000-3310 cell lines stably expressing a single chain antibody.

To confirm that the single cells identified by the model are representative of high-productivity CHO clonal cell lines, we separately transfected each of the 298 cell lines with DNA plasmids encoding either a single-chain antibody, monoclonal antibody, or Fc fusion bispecific antibody. Cell pools were generated by GS-based selection then pools evaluated in 14-day fed-batch culture to assess antibody productivity (**Figure 5b**). While numerous cell lines demonstrated higher-than-average pool titers for these antibodies, we identified subclone 1000-3310 as the lead candidate. Additional complex antibody sequences were produced to verify the applicability of this cell line as a platform production host (**Figure 5c**). Building on the success of this 43-day workflow, we assessed the kinetics of colony outgrowth for 1000-3310, as well as, this same cell line encoding and expressing an antibody following GS-selection (**Figure 5d**). The viable cell density of single cell-derived colonies peaked 14 days following deposition indicating that our outgrowth workflow could be reduced from 21 days to 14 days (**Figure 5a**). In addition to monoclonal origin, we were able to characterize baseline titer from clonal cell lines using an Octet RH96 before hitpick (**Figure 5e**). Utilizing this assay data as a binary readout for antibody production, we were able to further streamline this workflow and enrich for high-performing candidate cell lines.

To enhance our cell line engineering throughput, we scripted worklist-based liquid handling processes for a Lynx LM1800. Through the implementation of automated image ingestion, image review and hitpick list generation, automated cell culture passage and sampling, and Tetrascience-based vertical integration of instrument data, we achieved significant efficiency gains in cell line outgrowth leading to implementation of a 36-day end-to-end clonal cell line generation workflow.

## Discussion

This work presents an open-access workflow that automates image-based monoclonality verification and colony outgrowth quantification then seamlessly integrates the output into an automated cell culture workflow. Combining high-performance object detection with a robust colony segmentation method and optimized single cell outgrowth conditions, we define an end-to-end 36-day CLD process.

Among the object detection models evaluated for monoclonality verification, DETR demonstrated superior performance across both brightfield and fluorescent images. While the detection of single cells was consistently accurate (>0.9 F1-score), model performance was modestly lower for fluorescent images likely due to inherent features such as blurred boundaries (23) and light scattering (24). The misclassification of doublet and cell clusters in high-density training images are largely attributable annotation-related ambiguity. The structure of densely packed cells added subjectivity to manual labeling, particularly in instances where two adjacent doublets are classified together as a cluster.

For single cell images from a standard CLD workflow, DETR maintained high precision, recall, and F1 score. Fluorescent images generally present lower variation in cell morphology and image quality relative to brightfield images. While brightfield images initially showed lower scores at an IoU threshold of 0.5, relaxing the threshold to 0.25 or utilizing Dot Distance analysis (DotD), showed that detection centers were spatially accurate within 10 pixels. DotD analysis was applied to verify positional accuracy, defined as:

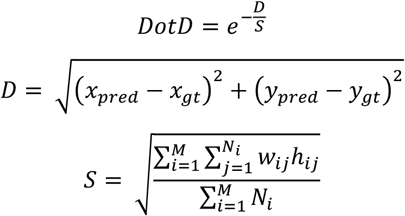

where (*x*_*pred*_, *y*_*pred*_)and (*x*_*gt*_, *y*_*gt*_)respectively denote the center coordinates of prediction and ground truth bounding boxes, *M* denotes the number of images in the dataset, *N*_*i*_ denotes the number of labeled bounding boxes in the *i*-th image, and *w*_*ij*_, *h*_*ij*_ denote the width and height of the *j*-th bounding box in the *i*-th image (25). DotD was initially developed for aerial image analysis where object size is small relative to image dimensions making this method well suited to evaluate single cell detection against the background of an image well. The formulation normalizes the center-to-center pixel distance *D* by the average object size and applies an exponential decay to produce a similarity score between 0 and 1, emphasizing positional alignment over box dimensions.

When the average object size *S* is estimated at 20 pixels, DETR precision, recall, and F1-score increased as the center-to-center threshold expanded achieving >0.9 near 10 pixels (**Figure 6a-c**). Precision rapidly rose as smaller thresholds eliminated false positives, while recall gradually improved as additional true positives were detected at larger thresholds. In CLD-relevant small-object detection, precise localization is often more important than mask overlap. Traditional IoU metrics can disproportionately penalize minor position errors,, whereas DotD provides a more task-relevant measure of positional accuracy.

**Figure 6.**
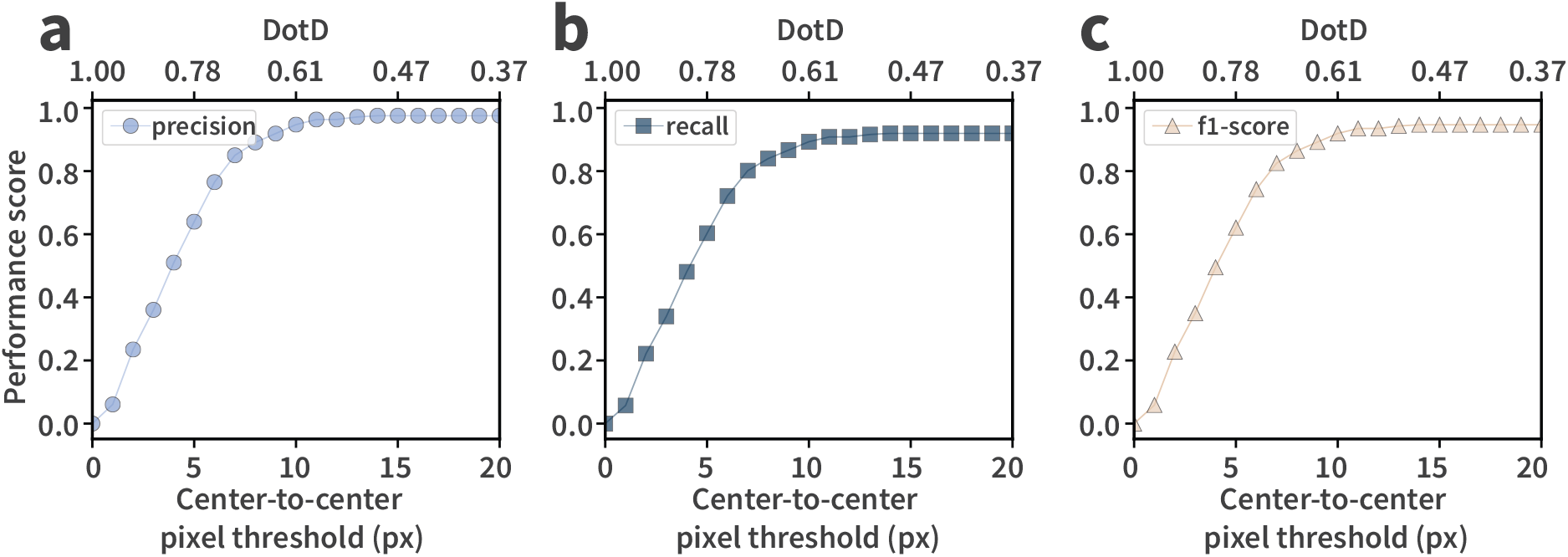
(a) Precision, (b) recall, and (c) F1-score as functions of center-to-center pixel distance threshold under the DotD metric. Scores increase with larger thresholds, reflecting improved tolerance for positional deviations. The top axis shows corresponding DotD similarity values, which decay exponentially from 1.00 to 0.37 as the threshold increases from 0 to 20 pixels. Around a 10-pixel threshold (DotD ≈ 0.61), all three metrics reach approximately 0.90 indicating strong detection performance at moderate positional tolerance.

We next compared feature-based segmentation with several zero-shot SAM2 architectures to quantify cell colony outgrowth. The feature-based method provided consistent estimates of confluence regardless of colony morphology. While the different SAM2 variants were effective for images of dense colonies, SAM2 struggled with sparse colonies lacking well-defined boundaries. The tiny and small architectures frequently under-segmented these colonies, while the base-plus and large models missed the colonies altogether. These limitations likely result from SAM2 reliance on hierarchical attention and mask generation, where prominent object boundaries are assumed. Similar challenges have been reported for SAM2 variants (26). Future improvements will explore Low-Rank Adaptation fine-tuning to enhance the sensitivity of SAM2 to sparse and low-contrast colonies while maintaining computational efficiency (27, 28).

Integration of CLAIRE output and a Lynx LM1800 liquid handler unified image-based decision-making with automated cell culture. Our ability to reduce cell line outgrowth from 21 to 14 days and automate hitpick list generation significantly increased daily throughput while reducing hands-on cell culture for error-prone activities like 96-well to 24-well hitpick. We validated this workflow by generating clonal cell lines capable of producing diverse antibodies at high titer.

## Conclusion

This study establishes a comprehensive workflow to automate cell line development campaigns through the integration of deep-learning image analysis and robotic liquid handling. We benchmarked multiple state-of-the-art object detection models and identified DETR as the highest performing architecture for cell monoclonality verification. Further, we show that feature-based detection outperforms several variants of SAM2 for colony confluence applications and highlights the need for specialized metrics such as DotD to more accurately quantify positional detection success for small objects relative to IoU thresholds. Collectively, the CLAIRE tool and Lynx LM1800 automation methods increase cell culture throughput and enable an end-to-end 36 day CLD workflow for pool to clone identification.

## Methods

### Cell Culture

CHOZN GS^-/-^ CHO cells were cultured in EX-CELL CD CHO Fusion Cell Culture Growth Medium supplemented with 3% (v/v) GlutaMAX. Suspension cell culture was performed at 37°C and 125 rpm in a humidified incubator with 5% CO_2_. Cells were passaged twice weekly, seeded first at 0.3E6 cells/mL for 3 day culture then seeded at 0.2E6 cells/mL for 4 day culture.

### Single Cell Deposition and Imaging

The viable cell density of CHOZN GS^-/-^ CHO cells was quantified using ViCell Blu then 1E6 cells collected by centrifugation (4 minutes, 4,000 rpm, room temperature). Spent medium was aspirated and the cells were resuspended in 1 mL PBS. One vial of CellTrace Violet dye was resuspended in 20 μl DMSO then 1 μl resuspended dye added to the cell suspension. Cells were incubated for 45 minutes at 37°C and 125 rpm in a humidified incubator with 5% CO_2_. Cells were collected by centrifugation (4 minutes, 4,000 rpm, room temperature), staining solution aspirated, and cells resuspended in 1 mL PBS.

Single cell cloning medium consisting of EX-CELL CHO Cloning Medium, 20% (v/v) filtered spent medium from CHOZN GS^-/-^ CHO cells, 2.5% (v/v) ClonaCell CHO ACF Supplement, and 3% (v/v) GlutaMAX was filtered through a 0.2 μm PES membrane. Next, 200 μl of filtered medium was added to each well of a 96-well plate and the plate incubated at 37°C until single cell deposition using an F.SIGHT 2.0. Following cell deposition, 10 μl of cell suspension was transferred to well C4 as a focus control then each 96-well plate was centrifuged (1 minute, 1,000 rpm, room temperature). A Celigo plate imager was used to acquire brightfield and fluorescence images for each well of each 96-well plate. Plates were incubated at 37°C for 9 or 14 days then, after colony outgrowth, a brightfield image of each well was captured. To each well of the 96-well plate, 100 μl EX-CELL CD CHO Fusion Cell Culture Growth Medium containing 3% (v/v) GlutaMAX was added and cells were triturated three times to disaggregate clumped cells. Images were processed for monoclonal occupancy and outgrowth to generate a hitpick list of clonal cell lines originating only from a single cell and colony with area >5% of the well.

### Manual Clonal Cell Line Generation Workflow

Clonal cell lines were hitpicked by transferring 200 μl of source cell suspension into 24-well deep-well plates containing 1 mL of warm EX-CELL CD CHO Fusion Cell Culture Growth Medium containing 3% (v/v) GlutaMAX. Cells were incubated 5 days at 37°C and 125 rpm in a humidified incubator with 5% CO_2_. The viable cell density of each clonal cell line was quantified using ViCell Blu then a volume corresponding to 0.3E6 cells transferred from the source 24-well plate into a destination 24-well deep-well plate containing (1.2 – volume of cells) mL warm EX-CELL CD CHO Fusion Cell Culture Growth Medium containing 3% (v/v) GlutaMAX. Cell passage in 24-well deep-well plates continued for three passages: first (3 days, 0.3E6 cells/mL in 1.2 mL), second (4 days, 0.3E6 cells/mL in 3 mL), and third (3 days, 0.3E6 cells/mL in 3mL). Clonal cell lines were scaled up using tubespin into 10 mL and 30 mL culture at 200 rpm and 250 rpm respectively. Cells were cryopreserved using a VIAFreeze Quad controlled rate freezer in EX-CELL CD CHO Fusion Cell Culture Growth Medium containing 3% (v/v) GlutaMAX and 7.5% (v/v) DMSO.

### Automated Clonal Cell Line Generation Workflow

Automation of the single cell cloning workflow was achieved using a Lynx LM1800 liquid handler following the 36-day workflow described in **Figure 5a**. The method wrapper *Day 0 Single Cell Print* automates addition of 200 μl medium to each well of barcoded 96-well plates without bubble creation. The method wrapper *Day 9 Colony Imaging* automates addition of 100 μl medium and dispersion of cell colonies. The method wrapper *Day 14 Hitpick* automates addition of 1.2 mL medium to each well of destination 24-well deep-well plates followed by resuspension of cells in the source 96-well plate in a five-point star pattern then list-based transfer of resuspended cells to destination 24-well deep-well plates. The method wrappers *Day 19 ViCell Blu Cell Count* and *Day 19 Passaging and Medium Exchange* automate cell culture sampling for ViCell Blu quantification, addition of variable volumes of medium to each well of destination 24-well deep-well plates, addition of variable volumes of source cell culture to achieve 0.3E6 cells/mL in 1.2 mL total volume, and aspiration of medium from centrifuged cell cultures below target density for resuspension and transfer into destination plates. The method wrappers *Day 22 ViCell Blu Cell Count, Day 22 Passaging*, and *Day 22 Titer and ddPCR Sampling* automate cell culture sampling for ViCell Blu quantification, addition of variable volumes of medium to each well of destination 24-well deep-well plates, addition of variable volumes of source cell culture to achieve 0.3E6 cells/mL in 3 mL total volume, and sample generation for copy number (cell pellet) and titer (supernatant) analyses. The method wrappers *Day 26 ViCell Blu Cell Count, Day 26 Passaging*, and *Day 26 NGS Sampling* automate cell culture sampling for ViCell Blu quantification, addition of variable volumes of medium to each well of destination 24-well deep-well plates, addition of variable volumes of source cell culture to achieve 0.3E6 cells/mL in 3 mL total volume, and sample generation for next-generation sequencing (NGS) assays. User input scripts with wrappers, base methods, and utility submethods are summarized in Supplemental Table 1.

### Dataset Preparation: Image Region of Interest Extraction and Annotation for Model Training

High-resolution images were acquired at 7530 × 7530 pixels. Individual cells occupied a spatial footprint of approximately 10-15 × 10-15 pixels and correspond to less than 0.0004% of the total image area. Training deep learning models on the entire image would be computationally intensive and inefficient due to the disproportionate size of the target object to the full image; therefore, regions of 256 × 256 pixels containing cells were extracted for modeling. Regions of interest were extracted from brightfield and fluorescent images then annotated using the *Make Sense* tool with labels exported in YOLO format for both object class and normalized bounding boxes. A total of 1,500 images were annotated into single cell, doublet, and cluster categories.

### Object Detection Model Training: YOLO-NAS

YOLO-NAS is an advanced single-stage detector built on the YOLO family then optimized using Neural Architecture Search technology (29). YOLO-NAS-L large variant was used for CLAIRE. Models were trained on custom datasets using the SuperGradients framework and utilized a cosine-annealing learning rate schedule (initial learning rate: 5e-4) with linear warmup for the first two epochs (start: 1e-4). Optimization used Adam with weight-decay 1e-4. Training ran for 100 epochs with batch size 3, as larger batch sizes led to non-convergence. Mixed precision was not used and checkpint averaging was applied for better generalization. The loss function was PPYoloELoss (use_static_assigner=False, reg_max=16) and model evaluation used DetectionMetrics_050 (score threshold: 0.1, top-k: 300). Post-processing was handled using PPYoloEPostPredictionCallback, configured with a score threshold of 0.01, non-maximum suppression top-k of 1000, maximum of 300 predictions, and non-maximum suppression threshold of 0.7. Performance was monitored using Mean Average Precision at IoU threshold 0.50 (mAP@0.50) with best checkpoint saved automatically.

### Object Detection Model Training: FRRF-v2

A Faster R-CNN (30) model incorporating a ResNet-50 backbone with a Feature Pyramid Network (31) was used to perform object detection on the annotated regions of interest. The model architecture was implemented using the Faster R-CNN ResNet50 FPN-2 v2 variant and pre-trained on the COCO dataset (weights: COCO-v1), selected for its balance between detection accuracy and computation efficiency. To adapt the model to the specific classification task, the final classification head was replaced with a FastRNNPredictor configured to output four classes: background, single cell, doublet, and cell cluster. Training and validation datasets were loaded using PyTorch’s DataLoader utility with a batch size of 4 for training and 2 for validation. Model training was conducted on a CUDA-enabled GPU using the Adam optimization algorithm with a learning rate of 0.001 and a weight decay coefficient of 0.0001. The training process was executed over a maximum of 100 epochs.

### Object Detection Model Training: DETR

The DETR model (32) was configured with a ResNet-50 backbone and utilized sine position embeddings with a transformer architecture comprising 6 encoder and 6 decoder layers, a hidden dimension of 256, 8 attention heads, and 100 object queries, while not employing dilation or instance segmentation masks. Training was performed on a custom dataset with 4 classes for 100 epochs on a CUDA-enabled GPU with a train batch size of 5, validation batch size of 2, a 1e-4 initial learning rate, a 1e-5 learning rate, and a 1e-4 weight decay while incorporating auxiliary losses and early stopping with a patience of 10. The loss function included class, Least Absolute Deviations (L1) for bounding box, and Generalized Intersection over Union costs weighted at 1, 5, and 2, respectively. An end-of-sequence (no object) coefficient of 0.1. The model was initialized using ‘detr-r50-e632da11.pth’ pre-trained weights.

### Object Detection Model Performance Evaluation

Model performance was assessed using a per-class framework that reports true positives, false positives, and false negatives. Predictions were first filtered using a confidence threshold of 0.7 and ranked by score to prioritize the most reliable detections. To avoid penalizing edge artifacts, any ground truth or predicted box within 15 pixels of the image boundary (roughly 6% of the 256 pixel image dimension) was excluded from evaluation. These exclusions occurred mainly in the prediction set because detection models often identify partially visible objects near image edges, whereas ground-truth annotation typically omitted cropped instances. Prediction and ground truth matching was performed using an IoU threshold of 0.5. Successful matches increased true positive counts and marked both boxes as matched. Any unmatched ground truth that was not ignored was counted as false negative, and unmatched predictions outside the boundary-ignore zone were counted as false positive. All objects were treated independently and nested relationships such as single cells within doublets or clusters were not considered. Performance metrics were computed as follows:

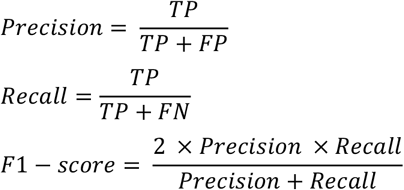

### Confluency Assessment: Feature-Based Workflow

Area of cell coverage within wells was quantified using a deterministic image-processing pipeline applied to brightfield images. Grayscale conversion was performed for color inputs followed by local contrast enhancement using CLAHE (clipLimit: 2.0, tileGridSize: 32 × 32). A coarse well mask was generated via intensity thresholding (value: 10) and Gaussian blurring (kernel: 5 × 5) was applied to suppress noise. Cellular texture was detected using the Laplacian operator (threshold: 10) to isolate textured regions. Morphological closing with an elliptical kernel (30 × 30) was used to merge fragmented detections into contiguous colonies. The colony mask was constrained to the well interior by intersecting with the largest external contour of the well mask. Confluency was calculated as the ratio of colony pixels to well pixels then expressed as a percentage.

### Confluency Assessment: Zero-Shot SAM2 Segmentation with Feature-Based Mask Refinement

Cell coverage was quantified using a zero-shot segmentation approach implemented with Segment Anything Model 2 (SAM2) (33). Brightfield images were standardized for inference by converting BGR to RGB and resizing to 1024 × 1024 pixels. Segmentation outputs were generated from all available hierarchical checkpoints: sam2.1_hiera_b+, sam2.1_hiera_s, sam2.1_hiera_t, and sam2.1_hiera_l. Inference settings were controlled through corresponding configuration .yaml files to ensure consistency. Automatic masks were produced using SAM2AutomaticMaskGenerator configured with parameters: points_per_side=32, points_per_batch=64, pred_iou_thresh=0.80, stability_score_thresh=0.95, min_mask_region_area=150 px, box_nms_thresh=0.7, output_multimask=False, crop_n_layers=1, and crop_overlap_ratio=0.0. Masks were refined through a post-processing screen in which regions located outside the well and areas affected by image-stitching artifacts were excluded. Confluency was expressed as the proportion of colony coverage relative to the well area.

### Lynx LM1800 Methods and CLAIRE User Interface

The Lynx LM1800 scripts and CLAIRE pre-trained models and user interface are available via https://github.com/Gilead-Public/CLAIRE. All model training used an NVIDIA RTX 5000 Ada Generation Laptop GPU and Windows operating system. Training workflows were implemented in Python (>3.9).

## References

1. ICH. Derivation and Characterisation of Cell Substrates Used for Production of Biotechnological/Biological Products Q5D. 1997.

2. FDA. Center for Biologics Evaluation and Research Food and Drug Administration Supplement to the Points to Consider in the Production and Testing of New Drugs and Biologic& Produced by Recombinant DNA Technology: Nucleic Acid Characterization and Genetic Stability. Food and Drug Administration (FDA) Pharmaceutical CGMPs. 1992.

3. WHO. Recommendations for the evaluation of animal cell cultures as substrates for the manufacture of biological medicinal products and for the characterization of cell banks. World Health Organization Technical Report Series, No 978. 2013.

4. Zhou X, Wu H, Wen H, Zheng B. Advances in Single-Cell Printing. Micromachines (Basel). 2022;13(1).

5. Scherzinger J, Türk D, Aprile-Garcia F. An optimized and validated workflow for developing stable producer cell lines with 99.99% assurance of clonality and high clone recovery. bioRxiv. 2022:2022.12.16.520697.

6. Steinwandter V, Borchert D, Herwig C. Data science tools and applications on the way to Pharma 4.0. Drug Discovery Today. 2019;24(9):1795–805.

7. O’shea K, Nash R. An introduction to convolutional neural networks. arXiv preprint 151108458. 2015.

8. Stirling DR, Swain-Bowden MJ, Lucas AM, Carpenter AE, Cimini BA, Goodman A. CellProfiler 4: improvements in speed, utility and usability. BMC Bioinformatics. 2021;22(1):433.

9. van Tol N, Rolloos M, van Loon P, van der Zaal BJ. MeioSeed: a CellProfiler-based program to count fluorescent seeds for crossover frequency analysis in Arabidopsis thaliana. Plant Methods. 2018;14(1):32.

10. Hennig H, Rees P, Blasi T, Kamentsky L, Hung J, Dao D, et al. An open-source solution for advanced imaging flow cytometry data analysis using machine learning. Methods. 2017;112:201–10.

11. Laan SNJ, Dirven RJ, Bürgisser PE, Eikenboom J, Bierings R, for the Sc. Automated segmentation and quantitative analysis of organelle morphology, localization and content using CellProfiler. PLOS ONE. 2023;18(6):e0278009.

12. Zhang H, Ericsson M, Virtanen M, Weström S, Wählby C, Vahlquist A, et al. Quantitative image analysis of protein expression and colocalisation in skin sections. Experimental Dermatology. 2018;27(2):196–9.

13. Gómez-de-Mariscal E, García-López-de-Haro C, Ouyang W, Donati L, Lundberg E, Unser M, et al. DeepImageJ: A user-friendly environment to run deep learning models in ImageJ. Nature Methods. 2021;18(10):1192–5.

14. Carpenter AE, Jones TR, Lamprecht MR, Clarke C, Kang IH, Friman O, et al. CellProfiler: image analysis software for identifying and quantifying cell phenotypes. Genome Biology. 2006;7(10):R100.

15. Fu X, Lin Y, Lin DM, Mechtersheimer D, Wang C, Ameen F, et al. BIDCell: Biologically-informed self-supervised learning for segmentation of subcellular spatial transcriptomics data. Nature Communications. 2024;15(1):509.

16. Stringer C, Wang T, Michaelos M, Pachitariu M. Cellpose: a generalist algorithm for cellular segmentation. Nature Methods. 2021;18(1):100–6.

17. Yapp C, Novikov E, Jang W-D, Vallius T, Chen Y-A, Cicconet M, et al. UnMICST: Deep learning with real augmentation for robust segmentation of highly multiplexed images of human tissues. Communications Biology. 2022;5(1):1263.

18. Archit A, Freckmann L, Nair S, Khalid N, Hilt P, Rajashekar V, et al. Segment Anything for Microscopy. Nature Methods. 2025;22(3):579–91.

19. Morelli R, Clissa L, Amici R, Cerri M, Hitrec T, Luppi M, et al. Automating cell counting in fluorescent microscopy through deep learning with c-ResUnet. Scientific Reports. 2021;11(1):22920.

20. Ferreira EKGD, Silveira GF. Classification and counting of cells in brightfield microscopy images: an application of convolutional neural networks. Scientific Reports. 2024;14(1):9031.

21. Schmidt U, Weigert M, Broaddus C, Myers G, editors. Cell detection with star-convex polygons. Medical image computing and computer assisted intervention–MICCAI 2018: 21st international conference, Granada, Spain, September 16-20, 2018, proceedings, part II 11; 2018: Springer.

22. Xu J, Wang A, Wang Y, Li J, Xu R, Shi H, et al. AICellCounter: A Machine Learning-Based Automated Cell Counting Tool Requiring Only One Image for Training. Neurosci Bull. 2023;39(1):83–8.

23. Haider SA, Cameron A, Siva P, Lui D, Shafiee MJ, Boroomand A, et al. Fluorescence microscopy image noise reduction using a stochastically-connected random field model. Scientific Reports. 2016;6(1):20640.

24. Cossarizza A, Chang HD, Radbruch A, Abrignani S, Addo R, Akdis M, et al. Guidelines for the use of flow cytometry and cell sorting in immunological studies (third edition). Eur J Immunol. 2021;51(12):2708–3145.

25. Xu C, Wang J, Yang W, Yu L. Dot Distance for Tiny Object Detection in Aerial Images. 2021 IEEE/CVF Conference on Computer Vision and Pattern Recognition Workshops (CVPRW). 2021:1192–201.

26. Pei J, Zhou Z, Zhang T. Evaluation study on SAM 2 for class-agnostic instance-level segmentation. arXiv preprint 240902567. 2024.

27. Mandal S, Karthikeyan D, Paldhe M. SAM2LoRA: Composite Loss-Guided, Parameter-Efficient Finetuning of SAM2 for Retinal Fundus Segmentation. arXiv preprint 251010288. 2025.

28. Forni B, Lombardi G, Pozzi F, Planamente M. FS-SAM2: Adapting Segment Anything Model 2 for Few-Shot Semantic Segmentation via Low-Rank Adaptation. arXiv preprint 250912105. 2025.

29. Research. YOLO-NAS A Next-Generation, Object Detection Foundational Model generated by Deci’s Neural Architecture Search Technology. Deci-AI 2023.

30. Ren S, He K, Girshick R, Sun J. Faster r-cnn: Towards real-time object detection with region proposal networks. Advances in neural information processing systems. 2015;28.

31. Li Y, Xie S, Chen X, Dollar P, He K, Girshick R. Benchmarking detection transfer learning with vision transformers. arXiv preprint 211111429. 2021.

32. Carion N, Massa F, Synnaeve G, Usunier N, Kirillov A, Zagoruyko S, editors. End-to-end object detection with transformers. European conference on computer vision; 2020: Springer.

33. Ravi N, Gabeur V, Hu Y-T, Hu R, Ryali C, Ma T, et al. Sam 2: Segment anything in images and videos. arXiv preprint 240800714. 2024.

